# PRMT5 in T helper lymphocytes is essential for cholesterol biosynthesis-mediated Th17 responses and autoimmunity

**DOI:** 10.1101/792788

**Authors:** Lindsay M. Webb, Shouvonik Sengupta, Claudia Edell, Stephanie A. Amici, Janiret Narvaez-Miranda, Austin Kennemer, Mireia Guerau-de-Arellano

**Affiliations:** School of Health and Rehabilitation Sciences, Division of Medical Laboratory Science, College of Medicine, Wexner Medical Center, The Ohio State University, Columbus, OH, USA; Institute for Behavioral Medicine Research, The Ohio State University, Columbus, OH, USA; Department of Microbial Infection and Immunity, The Ohio State University, Columbus, OH, USA; Department of Neuroscience, The Ohio State University, Columbus, OH, USA; Biomedical Sciences Graduate Program, The Ohio State University, Columbus, OH, USA

**Author notes:** Corresponding Author: Mireia Guerau-de-Arellano.

## Abstract

Protein Arginine Methyltransferase (PRMT) 5 catalyzes symmetric dimethylation of arginine, a post-translational modification involved in cancer and embryonic development. However, the role of PRMT5 in T helper (Th) cell polarization and Th cell-mediated disease has not yet been elucidated. Here we report that PRMT5 is necessary for Th17 differentiation and EAE, via enhancement of cholesterol biosynthesis and activation of ROR-γt. PRMT5 additionally controls thymic and peripheral homeostasis in the CD4 Th cell life cycle, as well as iNK T and CD8 T cell development or maintenance, respectively. Overall, our two conditional PRMT5 KO models that selectively delete PRMT5 in all T cells (T-PRMT5^Δ/Δ^) or Th cells (iCD4-PRMT5^Δ/Δ^) unveil a crucial role for PRMT5 in T cell proliferation, Th17 responses and disease. These results point to Th PRMT5 and its downstream cholesterol biosynthesis pathway as promising therapeutic targets in Th17-mediated diseases.

## Introduction

Arginine methylation is a post-translational modification of histones and other proteins that regulates important physiological processes, including embryonic development and stem cell biology. This modification is catalyzed by a group of ten Protein Arginine Methyl Transferase (PRMT) enzymes (1). Type I, II and III PRMTs catalyze asymmetric dimethylation (ADM), symmetric dimethylation (SDM) and monomethylation of arginine (1), respectively. While both PRMT5 and 9 can catalyze SDM, PRMT5 is considered responsible for the majority of SDM in the cell (2). A short and a long isoform of this protein can be produced from the *PRMT5* gene by alternative splicing. PRMT5 overexpression is common during cellular transformation and cancer. PRMT5 has been shown to methylate a number of targets, ranging from histones H4R3 and H3R8 to NF-κB and spliceosome proteins. PRMT5 thereby promotes cancer/stem cell proliferation and survival (3). High proliferation and NF-κB signaling are also common in immune cells such as B cells and T cells and PRMT5 has been shown to play an important role in B cell biology (4). A crucial role for PRMT5 in T cell responses and immune disease is also just beginning to be recognized.

We recently reported that PRMT5 undergoes induction after T cell activation in a process controlled by NF-κB/MYC/mTOR signaling (5, 6). PRMT5’s SDM mark has also been shown to be dynamically regulated in T cells (7), suggesting it plays important functions. Indeed, we showed that both PRMT5 inhibitors and sh-RNA genetic knockdown impair T cell proliferation (6). Subsequently, genetic deletion of one of the PRMT5 isoforms in all T cells recapitulated this proliferation defect (8). However, it is not known how dual-isoform PRMT5 deletion impacts T cell proliferation and differentiation of Th cells into Th1/Th2/Th17/Treg phenotypes of importance in autoimmune and other diseases. Inflammatory Th1 and Th17 responses are particularly pathogenic, as they drive chronic tissue damage in autoimmune diseases such as Multiple Sclerosis (MS) (9).

Metabolic reprogramming upon T cell activation is a phenomenon that is increasingly recognized as an essential part of regulating Th cell function and polarization. Activated T cells grow and proliferate very rapidly, requiring the induction of a biosynthetic phenotype. Thus, quiescent naive or resting memory T cells, which rely on oxidative phosphorylation and/or fatty acid oxidation for energy generation, rapidly shift upon activation to biosynthetic metabolic pathways including glycolysis and cholesterol biosynthesis. Inflammatory Th1 and Th17 cells, in contrast to Tregs, require this highly glycolytic and biosynthetic reprogramming and cholesterol biosynthesis is particularly important for Th17 cell differentiation (10). However, the driver/s promoting the metabolic activation of the Th17 program is/are unknown.

Here we find that the arginine methyltransferase PRMT5 is required for Th17 differentiation via enhancement of cholesterol biosynthesis and activation of ROR-γt. Consequently, PRMT5 expression in Th cells was required for the Th17-mediated disease EAE. PRMT5-catalyzed methylation could influence multiple layers of the Th cell life cycle, from thymic development to peripheral maintenance and Th differentiation. Indeed, we find that PRMT5 controls all three of these processes, with strong impacts on CD4 Th cells at the thymic, peripheral homeostasis and polarization cell stages. We also observe important defects in iNK T and CD8 T cell development or maintenance, respectively, in the absence of PRMT5. Overall, our data unveil a crucial role for PRMT5 in T cell proliferation, Th17 responses and disease and point to Th PRMT5 and its downstream cholesterol biosynthesis pathway as promising therapeutic targets in Th17-mediated diseases.

## Results

### Development of constitutive pan-T cell-specific (T-PRMT5^Δ/Δ^) and inducible CD4 Th cell-specific (iCD4-PRMT5^Δ/Δ^) mouse models of PRMT5 deficiency

PRMT5 is essential for embryonic development (11, 12) and hematopoietic cells development (13). Therefore, evaluation of PRMT5’s function in T cells requires conditional KO models that allow PRMT5 deletion in a T cell subset-specific and time-controlled manner. To develop conditional PRMT5 KO mice where both PRMT5 protein-coding isoforms (**Fig. 1A**) are specifically deleted in T cells, we used the *Prmt5*^tm2c(EUCOMM)wtsi^ mutation that flanks exon 7 with loxP sites (13). To delete PRMT5 in all T cells (pan-T) or in the CD4 Th compartment, the PRMT5^fl/fl^ mice were crossed to CD4-cre (14) or CD4-cre-ERT2 (15) mice, respectively. The CD4-cre transgene is constitutively expressed in all CD4 expressing cells, which would include the thymic double positive T cells which would later develop into all CD3^+^ T cells, thereby providing a mouse model in which all peripheral T cells lack PRMT5 (**Fig. 1B**, T-PRMT5^Δ/Δ^ mice). In contrast, the tamoxifen-inducible CD4-cre-ERT2 transgene majorly provides peripheral CD4 T cell-specific deletion upon tamoxifen treatment (**Fig. 1B**, iCD4-PRMT5^Δ/Δ^ mice) with a lesser impact on DP thymocytes present during the tamoxifen treatment window (15). As expected, the short PCR product corresponding to *Prmt5* KO was amplified from both CD4^+^ Th and CD8^+^ Tc cells in T-PRMT5^Δ/Δ^ mice (**Fig. 1C**) but only in CD4^+^ Th cells from tamoxifen-treated iCD4-PRMT5^Δ/Δ^ mice (**Fig. 1D**).

**Figure 1.**
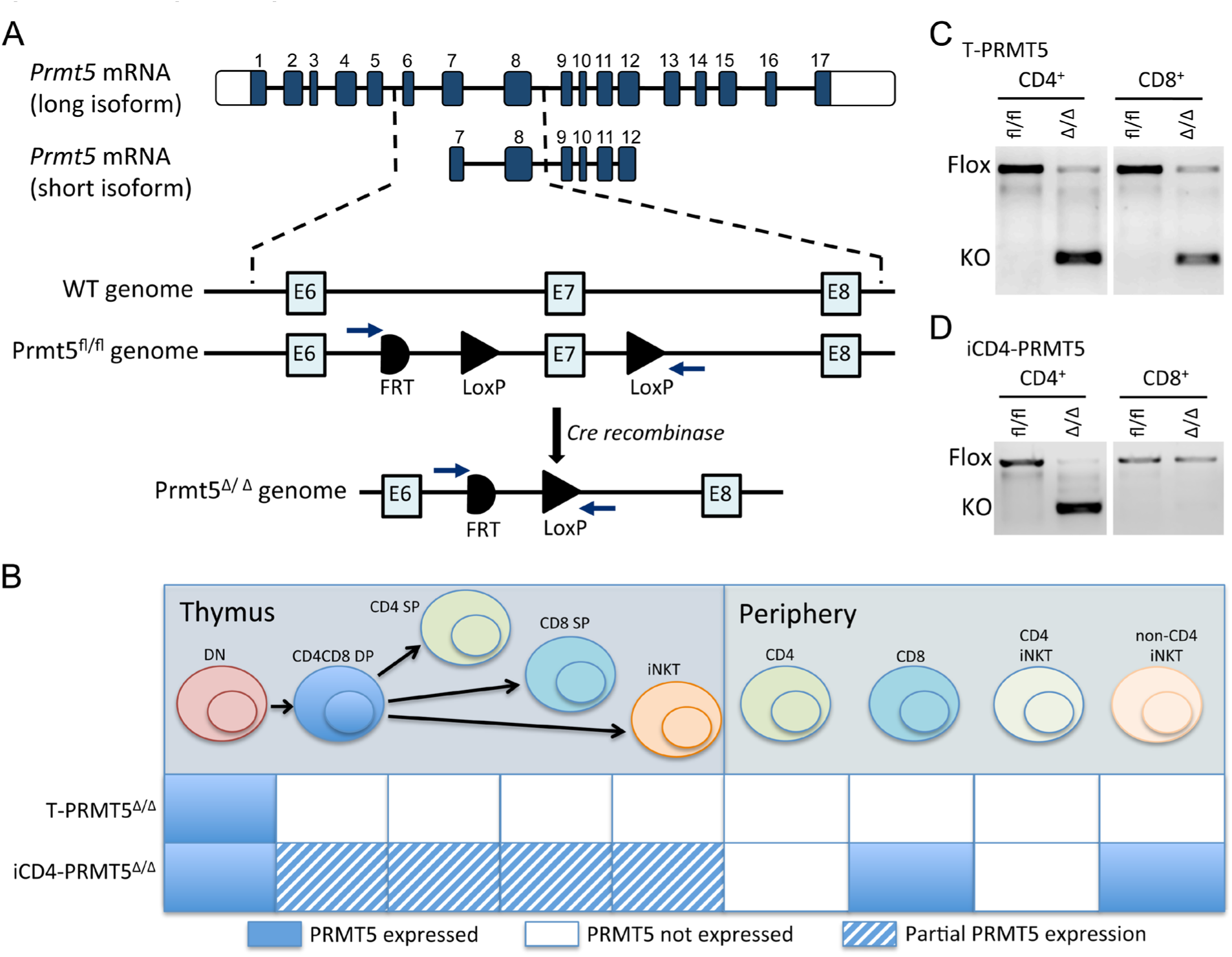
Constitutive and inducible CD4^+^ PRMT5 knockout models. (**A**) Schematic of *Prmt5* long and short isoform transcripts and *Prmt5* genomic locus targeting strategy. Exon 7, a common exon to all protein-coding *Prmt5* isoforms, was flanked by loxP sites to provide deletion of both isoforms in cre-expressing cells. (**B**) Schematic of expected PRMT5 deletion in thymus T cell precursors and peripheral T cells in two different transgenic models, namely T-PRMT5^Δ/Δ^ and iCD4-PRMT5^Δ/Δ^. (**C-D**) PCR amplification of genomic DNA isolated from CD4^+^ and CD8^+^ T cells from (**C**) T-PRMT5^Δ/Δ^ or (**D**) iCD4-PRMT5^Δ/Δ^ mice, as well as control PRMT5^fl/fl^ mice. PRMT5 KO band: 283bp; full length floxed PRMT5 band: 1100bp.

### PRMT5 is necessary for normal thymic CD4 Th cell and regulatory T cell (Treg) development

We have previously shown that PRMT5 is essential for T cell proliferation (6). Since thymocyte proliferation is a crucial event during T cell development, we evaluated the impact of PRMT5 deficiency on the thymic immune compartment by flow cytometry (**Fig. 2A**). Total thymic numbers were slightly but not significantly reduced in T-PRMT5^Δ/Δ^ mice compared to PRMT5^fl/fl^ and CD4-cre controls (**Fig. 2B**). The CD4-cre driver is first expressed at the DP stage. As expected, while total DN thymocyte numbers were unaffected (**Fig. 2C**), a significant loss in the DP compartment (**Fig. 2D**) was present. This defect was more prominent in the CD4SP (**Fig. 2E**) and Treg (**Fig. 2F**) populations. Interestingly, the loss of CD4SP cells was also observable in heterozygous PRMT5 KO (**Fig. 2E**), indicative of PRMT5 haploinsufficiency during thymic CD4 T cell development. In contrast, the CD8SP compartment was not significantly affected (**Fig. 2G)**. Consistent with the role of PRMT5 in proliferation, the observed thymic defects appear to be due to reduced thymocyte expansion, as no significant impact on DN, DP or SP frequencies was observed with the exception of reduced Treg frequency (**Supplemental Fig 1**).

**Figure 2.**
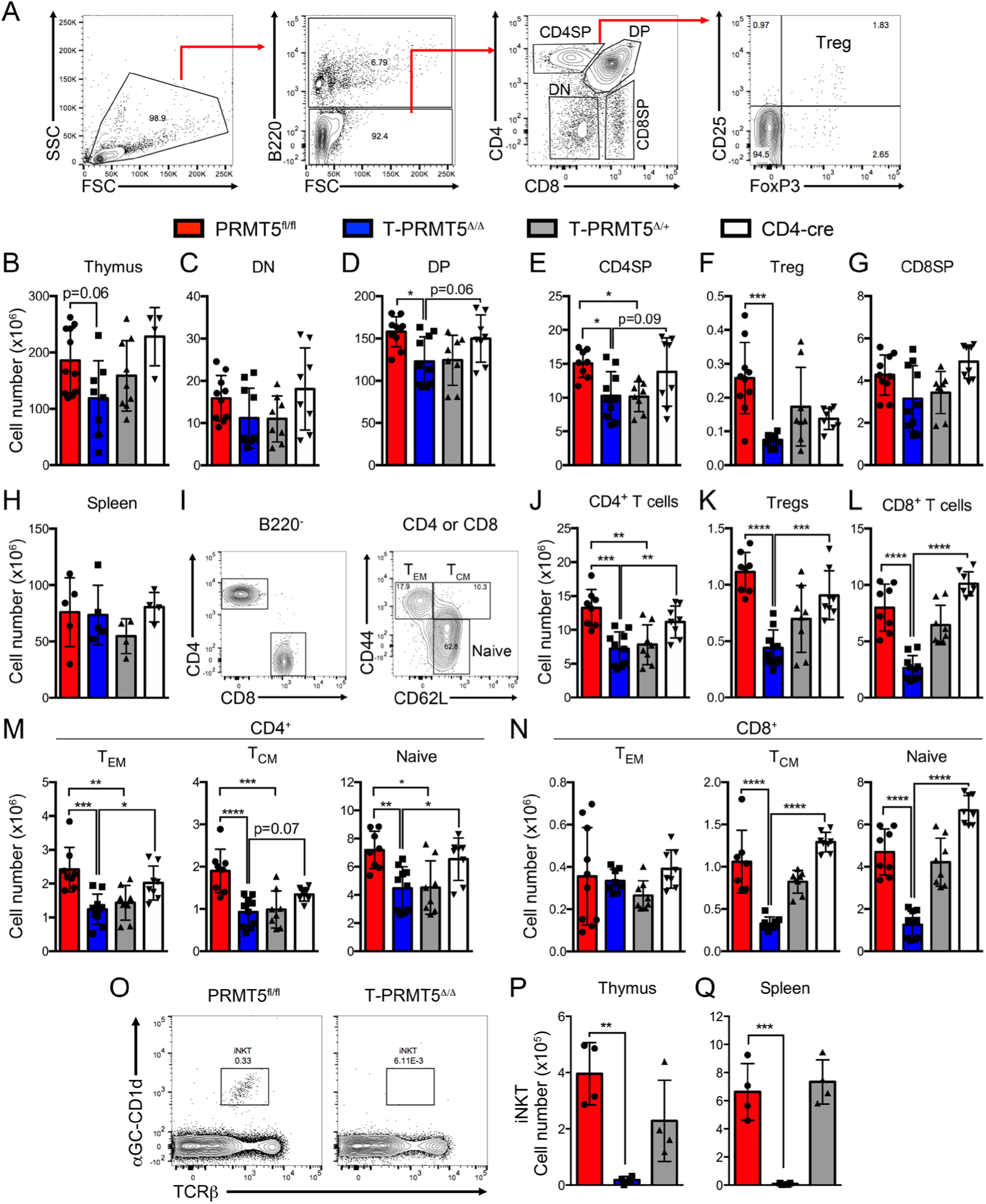
Impaired thymocyte development and peripheral T cell compartment in T-PRMT5^Δ/Δ^ mice. (**A**) Gating strategy for analysis of thymocyte populations. Live/dead gating was also performed after FSS/SSC but is not displayed. (**B-G**) Thymocytes were analyzed by flow cytometry and (**B**) total, (**C**) DN, (**D**) DP, (**E**) CD4SP, (**F**) Treg, and (**G**) CD8SP thymocyte numbers were calculated. (**H-N**) Splenocytes were analyzed by flow cytometry (gating strategy in **I**) and (**H**) total splenocyte, (**J**) CD4^+^, (**K**) Treg, (**L**) CD8^+^, (**M**) CD4^+^ T_EM_, T_CM_, and naive, and (**N**) CD8^+^ T_EM_, T_CM_, and naïve populations were calculated. (**O**) iNK T cell flow cytometric analysis gating strategy and total (**P**) thymic and (**Q**) splenic iNKT cell numbers. Data are representative of four independent experiments. Shown n = 5 independent mice of matched age. One-way ANOVA, followed by Dunnett’s multiple comparison test: *p<0.05, **p<0.01, ***p<0.001, ****p<0.0001.

To evaluate the impact of PRMT5 deletion on peripheral immune populations, we analyzed spleen and lymph nodes. While total splenocyte and lymph node cell numbers were unaffected in T-PRMT5^Δ/Δ^ mice (**Fig. 2H** and data not shown), flow cytometry analyses of T cell populations (**Fig. 2I**) showed reduced numbers of CD4 Th cells and Tregs in the spleen (**Fig. 2J-K**) and LNs (data not shown), following the same pattern observed in the thymus. This suggests that CD4 T cell defects could stem from thymic defects alone or, alternatively, combined thymic/peripheral homeostasis defects. In contrast, thymic CD8 SP numbers were normal, but a robust loss of CD8^+^ Tc cells was evident in T-PRMT5^Δ/Δ^ mice spleen (**Fig. 2L**) and LNs (data not shown), suggesting that CD8 T cell maintenance requires PRMT5. Among CD4 Th cells, all naive (CD62L^+^CD44^−^), effector memory (T_EM,_ CD62L^−^CD44^+^ effector), and central memory (T_CM_, CD62L^+^CD44^+^) CD4^+^ T cells were reduced (**Fig. 2M**). Within the CD8 T cell compartment, both the naive, and T_CM_ CD8^+^ T cells were drastically lost while no changes in T_EM_ cells were observed (**Fig. 2N**). Finally, as previously reported (8), we observed complete loss of iNKT cell populations in both thymus and spleen of T-PRMT5^Δ/Δ^ mice (**Fig. 2O-Q**). Overall, these data show that PRMT5 is required for thymic CD4 Th, Treg and iNK T cell development and normal CD4, CD8 and iNKT peripheral T cell compartments and that these defects are T cell intrinsic.

The observation that CD8^+^ T cell numbers were normal at the thymic level but robustly reduced peripherally suggests that CD8^+^ T cells are highly dependent on PRMT5 in the periphery. The peripheral CD4^+^ Th cell loss could result from thymic development defects or and/or peripheral homeostasis defects. To address this, we took advantage of the iCD4-PRMT5^Δ/Δ^ model, which allows temporally-controlled PRMT5 deletion, mostly impacting peripheral CD4 Th cells. To rule out any effects on thymic development, we evaluated thymocytes and peripheral immune cell compartments in adult iCD4-PRMT5^Δ/Δ^ mice (**Supplemental Fig. 2A**). As expected, thymic compartments were unaffected in iCD4PRMT5^Δ/Δ^ mice after 1 week of tamoxifen treatment (**Supplemental Fig. 2B-M**), indicating normal T cell development in these mice. We observed no significant effects of acute PRMT5 deficiency on peripheral CD4, CD8 and iNK T cell compartments (**Supplemental Fig. 2N-Y**). Extended 5-week tamoxifen-induced cre activity (**Supplemental Fig. 3A**) did not substantially impact thymic CD4SP, but instead reduced CD4 T cell numbers in the spleen (**Supplemental Fig. 3B-I**) and LNs (data not shown), confirming that PRMT5 promotes peripheral CD4 T cell homeostasis. Among peripheral CD4 T cells, both naive and T_EM_ subsets were lost (**Supplemental Fig. 3J**). We confirmed that this model is selective for CD4^+^ T cells, as CD8^+^ Tc populations were unaffected with extended tamoxifen treatment (**Supplemental Fig. 3F, I**). In summary, these data demonstrate that PRMT5 expression promotes both thymic T cell development and peripheral CD4 and CD8 T cell maintenance.

### PRMT5 drives CD4^+^ Th cell proliferation

To evaluate the impact of PRMT5 deficiency on peripheral Th cell function, we isolated CD4^+^ Th cells from T-PRMT5^Δ/Δ^ and analyzed them 48 hours after TcR engagement. We confirmed the near complete loss of PRMT5 protein expression in T-PRMT5^Δ/Δ^ Th cells compared to controls (**Fig. 3A, B**). The type I methyltransferase PRMT1 was also slightly reduced (**Fig. 3A, C**), confirming previously reported positive modulation of PRMT1 by PRMT5 (6). Functionally, such PRMT5 loss resulted in a robust (≥ 80%) suppression of proliferation in both CD4 Th cells (**Fig. 3D**) and total CD3^+^ T cells (**Supplemental Fig. 4**). To determine whether the proliferative defect is also present in T cells deleted of PRMT5 after thymic development, we analyzed Th cells from iCD4PRMT5^Δ/Δ^ mice. We found that PRMT5 protein induction after TcR-stimulation was suppressed to a lesser extent in this model, approximately 60-70% (**Fig. 3E, F**), consistent with the lower efficiency of the iCD4-cre-ER driver (15). PRMT1 maintained normal expression (**Fig. 3E, G**). In this model, Th cell proliferation was again suppressed (**Fig. 3H**). These data conclusively demonstrate a driver role for PRMT5 in TcR-induced CD4 Th and CD8 Tc cell proliferation.

**Figure 3.**
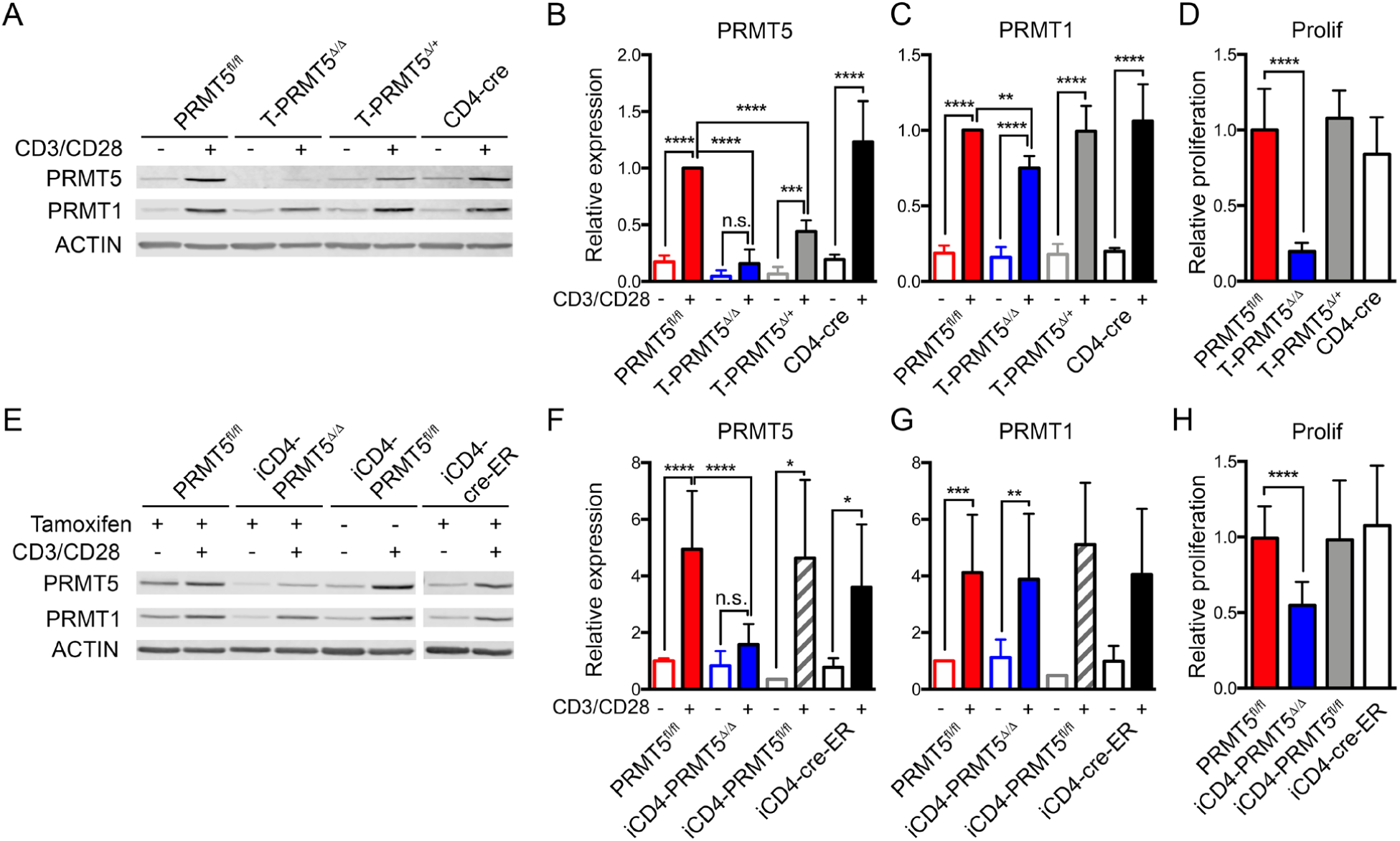
*Prmt5* deficiency suppresses T cell proliferation. Whole CD4^+^ T cells from (**A-D**) T-PRMT5^Δ/Δ^ or (**E-H**) iCD4-PRMT5^Δ/Δ^ mice were isolated, cells were collected for western blotting directly *ex vivo* and after 48 hours of anti-CD3/CD28 activation, and (**A, E**) analyzed by Western blot. Quantification of Western blot **(B, F)** PRMT5 and (**C, G**) PRMT1 expression. Data are pooled from at least three representative independent experiments (n=5-7 independent mice). (**D, H**) Proliferation of CD4^+^ T cells was analyzed by ^3^H-thymidine incorporation and expressed as a relative proliferation ratio to the resting PRMT5^fl/fl^ control condition. Data include at least three independent experiments (n=5-12 mice/group). One-way ANOVA, followed by Dunnett’s multiple comparison test. *p<0.05, **p<0.01, ***p<0.001, ****p<0.0001.

### PRMT5 regulates Th cell differentiation

Naive Th cell activation leads to differentiation into various Th cell phenotypes depending on the environmental milieu. Inflammatory Th1 and Th17 responses are highly pathogenic and drive chronic tissue damage in the autoimmune disease of the central nervous system MS (9). We first evaluated the impact of PRMT5 on Th cell differentiation in the T-PRMT5^Δ/Δ^ model where T cells lose PRMT5 during thymic development (**Fig. 4A**). Surprisingly, we observed increased Th1 differentiation among live CD44^+^ cells, following a slight defect in IFN-γ-secreting Th1 cells at day 3 (see **Fig. 4B-E**). These results are in contrast to the reduced Th1 responses observed in mice treated with PRMT5 inhibitors (6). Th17 differentiation was instead severely blunted, with abrogation of RORγt^+^IL-17^+^ Th17 cells, IL-17 secretion and RORγt expression (**Fig. 4F-I**). To sort out whether these effects are mediated by PRMT5-controlled changes at the thymic development vs. peripheral polarization steps, we used the iCD4-PRMT5^Δ/Δ^ model (**Fig. 4J**). In addition, approximately 50% of normal T cell proliferation remains upon TcR engagement in the iCD4-PRMT5^Δ/Δ^ model, ensuring a larger T cell pool. Th1 polarization was again increased with PRMT5 deficiency in this model, with the exception of reduced IFN-γ secretion at day 3, a defect recovered by day 7 (**Fig. 4K-N**). Likewise, Th17 polarization was practically absent, with lack of IL-17 secretion and RORγt induction (**Fig. 4O-R**). Overall, these data indicate that PRMT5 suppresses Th1 differentiation but is essential for Th17 differentiation.

**Figure 4.**
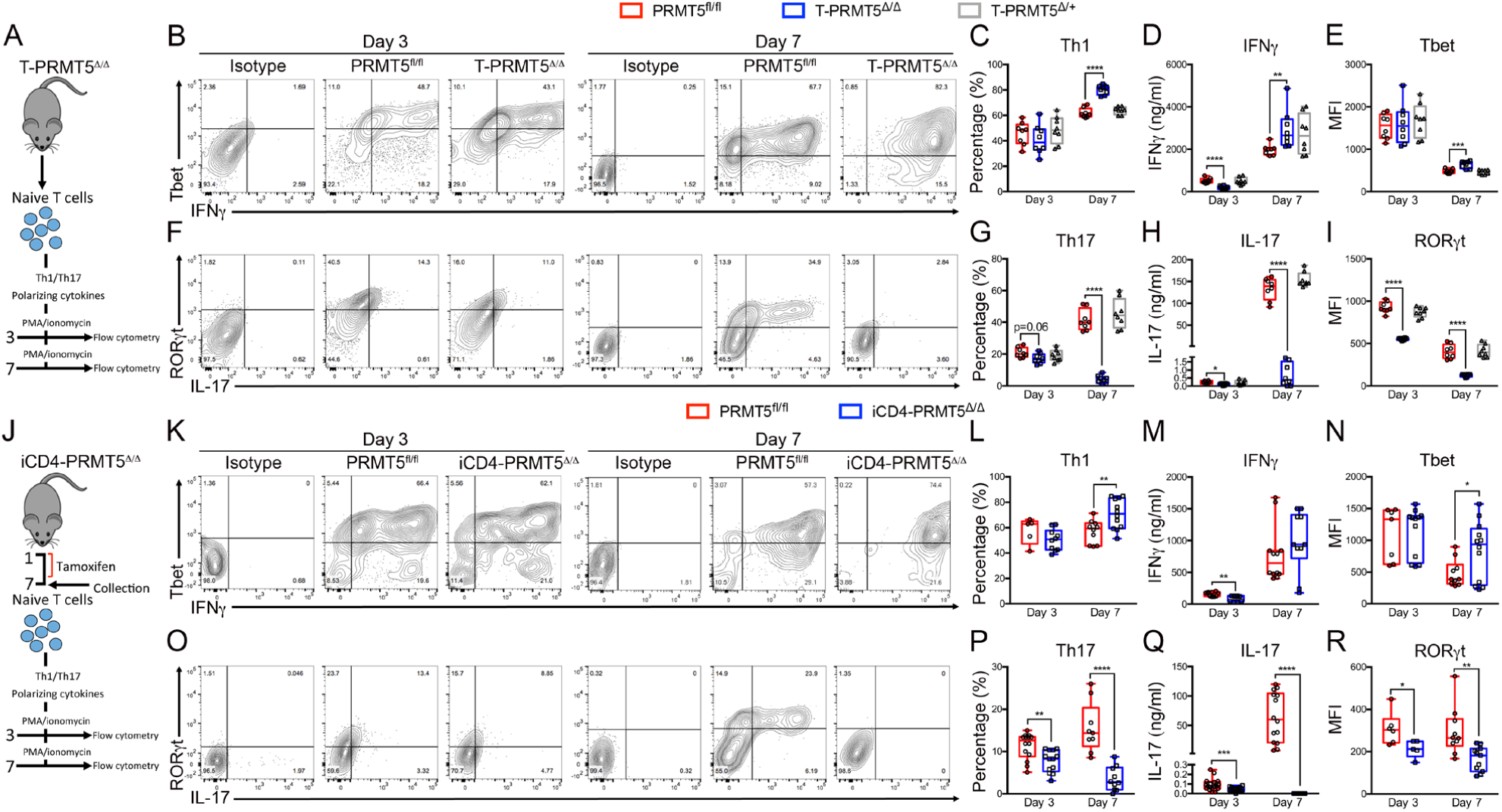
*Prmt5* deficiency abrogates Th17 cell differentiation. Naive CD4^+^ T cells isolated from (**A-I**) T-PRMT5^Δ/Δ^ or (**J-R**) iCD4-PRMT5^Δ/Δ^ mice were polarized into (**B-E, K-N**) Th1 or (**F-I, O-R**) Th17, and assessed by flow cytometry (**A** and **J** show experimental design schematic). Cells shown are gated on live CD44^+^ cells. Th1 cells were assessed by (**C, L**) Tbet^+^IFNγ^+^ cell %, (**D, M**) ELISA-analyzed IFNγ in cell supernatants, and (**E, N**) T-bet mean fluorescence intensity (MFI) by flow cytometry. Th17s were assessed by (**G, P**) RORγt^+^IL-17^+^ cell %, (**H, Q**) supernatant IL-17 detection by ELISA, and (**I, R**) RORγt MFI by flow cytometry. Data are pooled from 4-6 independent mice. One-way ANOVA, followed by Dunnett’s multiple comparison test, or Student’s t test, were used as appropriate. *p<0.05, **p<0.01, ***p<0.001, ****p<0.0001.

### PRMT5 regulates cholesterol biosynthesis to drive Th17 differentiation

Our data show that PRMT5 deficiency in T cells robustly suppresses Th cell proliferation and Th17 differentiation. To define the mechanisms by which PRMT5 achieves these effects, we performed RNA sequencing (RNAseq) in resting and activated CD4 Th cells from iCD4-PRMT5^Δ/Δ^ and control PRMT5^fl/fl^ mice. Lack of PRMT5 resulted in reduced expression of genes involved in lipid metabolism pathways, as revealed by Ingenuity Pathway Analysis (**Fig. 5A**). In particular, expression of multiple cholesterol biosynthesis pathway enzymes (yellow ovals in **Fig. 5B** pathway) was suppressed in PRMT5 deficient Th cells (**Fig. 5C**). The loss of several of these enzymes, namely *Tm7sf2, Hmgcs1, Lss* and *Acat2*, was validated by Real-Time PCR in cells undergoing Th17 differentiation (**Fig. 5D**). Lipid metabolism and cholesterol biosynthesis is not only essential for T cell growth and division (16-19), but also crucial for Th17 cell differentiation (20-22). Several cholesterol biosynthesis pathways intermediates, including lanosterol, zymosterol and desmosterol, are strong RORγ agonists essential for Th17 cell differentiation (20) and are highlighted in blue in the pathway (**Fig. 5B**). This led us to hypothesize that PRMT5 promotes Th17 differentiation by promoting production of cholesterol biosynthesis intermediates. If PRMT5 mediates the Th17 differentiation defect through suppression of cholesterol precursor biosynthesis, restoring such precursors in PRMT5 KO T cells should restore this defect. To test this, we supplemented cholesterol intermediates lanosterol or desmosterol during Th17 differentiation in iCD4-PRMT5^Δ/Δ^ T cells. We found that desmosterol enhanced Th17 differentiation in PRMT5^fl/fl^ T cells and restored normal levels of ROR-γt^+^IL-17^+^ (**Fig. 5E**) and IL-17^+^ (**Fig. 5F**) Th17 cell differentiation in iCD4-PRMT5^Δ/Δ^ T cells. In contrast, the further upstream lanosterol intermediate had no effect (**Fig. 5E-F**). Overall, these data show that PRMT5 promotes Th17 cell differentiation by enhancing cholesterol intermediate biosynthesis.

**Figure 5.**
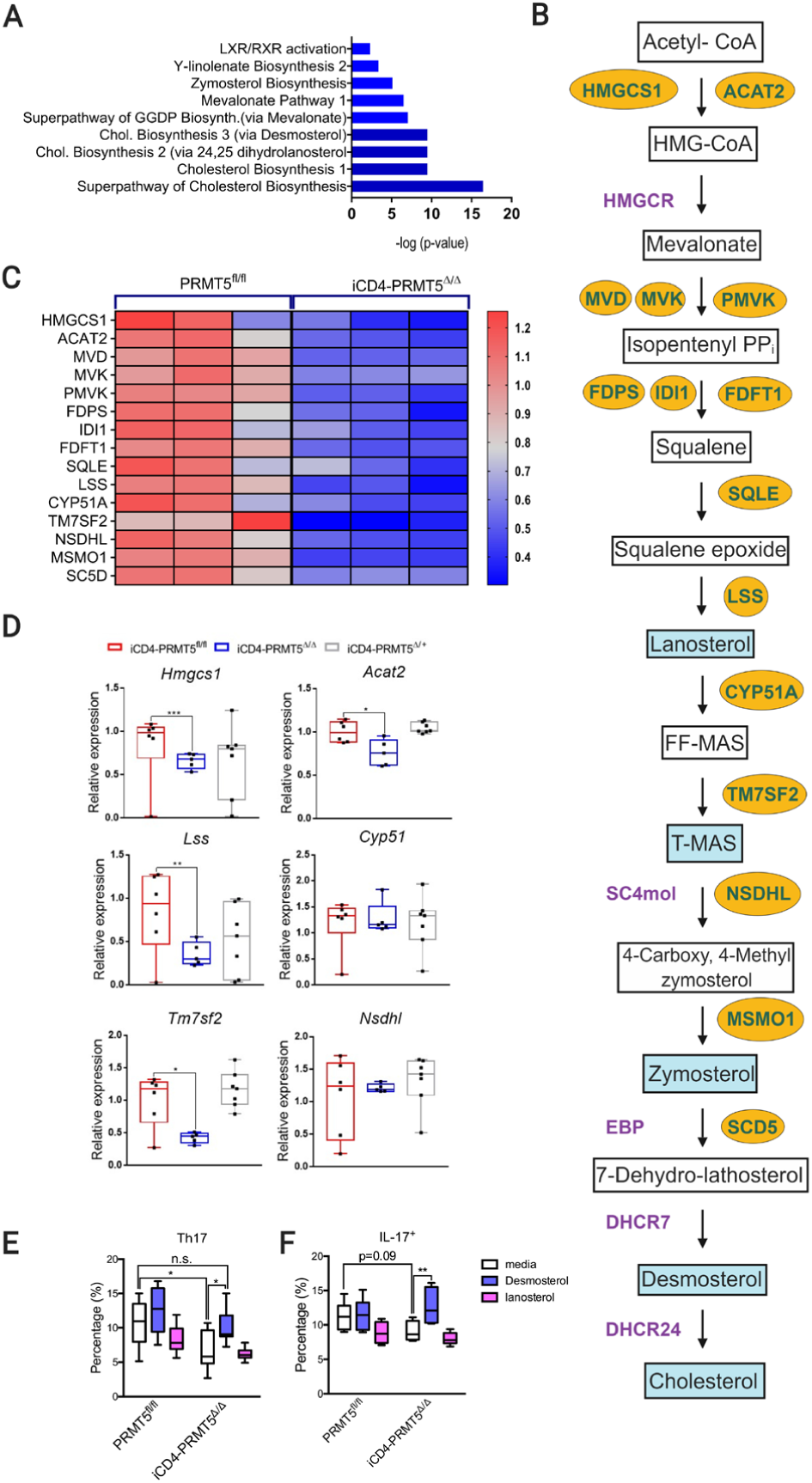
PRMT5 promotes cholesterol biosynthesis to drive Th17 differentiation. (**A**) Ingenuity pathway analysis of RNAseq gene expression of the top 9 down-regulated pathways in CD3/CD28-activated CD4 T cells from iCD4-PRMT5^Δ/Δ^ vs. iCD4-PRMT5^fl/fl^ mice. (**B**) Pathway of cholesterol biosynthesis. Enzymes differentially decreased in iCD4-PRMT5^Δ/Δ^ T cells are shown in yellow ovals. (**C**) Heatmap of cholesterol biosynthesis enzymes differentially expressed in iCD4-PRMT5^Δ/Δ^ mice and control iCD4-PRMT5^fl/fl^ mice CD4^+^ T cells. Each column represents a biological replicate. Up-regulated genes are shown in red and down-regulated genes are shown in blue. (**D**) Real time PCR of cholesterol biosynthesis pathway enzyme expression 3 days post-activation of naive CD4^+^ T cells of PRMT5^fl/fl^, iCD4-PRMT5^Δ/Δ^ or iCD4-PRMT5^Δ/+^ mice in Th17-polarizing conditions. (**E**) iCD4-PRMT5^Δ/Δ^ naive CD4^+^ T cells were isolated and activated in Th17 polarizing conditions in the presence or absence of cholesterol precursors desmosterol or lanosterol. Flow cytometric analysis of RORγt^+^IL-17^+^ or IL-17^+^ (gated on live CD4^+^CD44^+^) Th17 cells was performed 3 days post-activation. Data are pooled from two independent experiments (n=5). Two-way ANOVA, followed by Dunnett’s multiple comparison test. *p<0.05, **p<0.01, ***p<0.001, ****p<0.0001.

### PRMT5 is necessary to drive CD4^+^ Th cell pathogenesis and EAE autoimmunity

Th17 responses drive several autoimmune and inflammatory diseases, including MS. Given the crucial role of PRMT5 in Th17 differentiation, we hypothesized that loss of PRMT5 in the Th cell compartment would suppress EAE autoimmunity.

To test this hypothesis, we used the CD4 Th cell-specific (**Fig. 6A**) iCD4-PRMT5^Δ/Δ^ PRMT5 KO model. EAE was completely abolished in iCD4-PRMT5^Δ/Δ^ mice while mice with a heterozygous deletion of PRMT5 in Th cells experienced delayed mild disease (**Fig. 6B**). Disease development in PRMT5^fl/fl^ mice was associated with significant weight loss, while iCD4-PRMT5^Δ/+^ and iCD4-PRMT5^Δ/Δ^ maintained their weight (**Supplemental Fig. 5A**). Total CNS infiltrating T cell numbers were robustly reduced in iCD4-PRMT5^Δ/Δ^ mice (**Fig. 6C**), even as normal T cell numbers exist in the periphery (**Supplemental Fig. 2T-Y**). Within CD4 Th cells, important losses were evident in the CD44^+^CD62L^−^ T_EM_ and CD44^+^CD62L^+^ T_CM_ Th cells compartments (**Fig. 6D-E**). This indicates that Th-specific PRMT5 deficiency in Th cells is sufficient to impair recruitment of memory Th cells, presumably with pathogenic phenotypes, into the CNS. To address this, myelin-specific T cell proliferation and pathogenic Th1 and Th17 responses were evaluated in the CNS. Infiltrating CNS cells from iCD4-PRMT5^Δ/Δ^ mice did not proliferate (**Fig. 6F**) or secrete IFNγ (**Fig. 6G**) or IL-17 (**Fig. 6H**) in response to MOG. In particular, a substantial loss of MOG-specific Tbet^+^IFNγ^+^ Th1 cells (**Fig. 6I**), RORγt^+^IL-17^+^ Th17 cells (**Fig. 6J**), and the particularly pathogenic Tbet^+^IL-17^+^ Th17 population (**Fig. 6K**) was observed.

**Figure 6.**
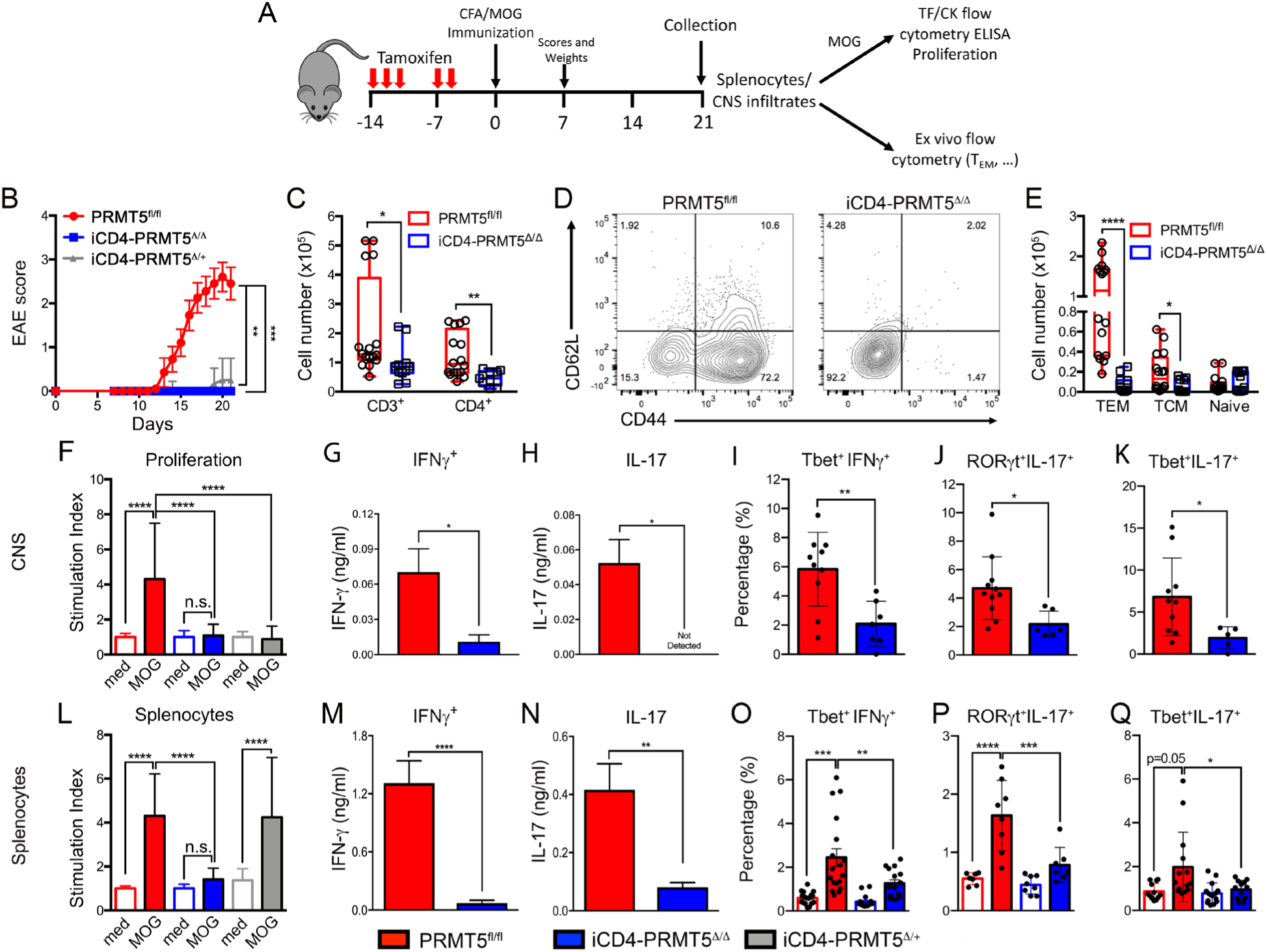
Th cell specific *Prmt5* deficiency prevents induction of EAE autoimmunity. (**A**) Schematic of tamoxifen treatment/EAE experimental design and downstream analyses. (**B**) EAE score in iCD4-PRMT5^Δ/Δ^ and indicated controls after MOG_35-55_/CFA immunization. All mice were treated with tamoxifen by oral gavage prior to immunization. (**C-E**) Flow analysis and quantification of CNS infiltrating (**C**) CD3^+^ and CD3^+^CD4^+^ T cells or (**D-E**) naive, T_EM_ and T_CM_ phenotype CD4 T cells from iCD4-PRMT5^Δ/Δ^ and indicated controls 21 days post-MOG_35-55_/CFA immunization. (**F-Q**) Infiltrating CNS cells (**F-K**) or splenocytes (**L-Q**) from day 14 iCD4-PRMT5^Δ/Δ^ and indicated controls were reactivated with MOG_35-55_. (**F, L**) Proliferation was monitored by ^3^H-thymidine incorporation and expressed as a relative proliferation ratio to the resting PRMT5^fl/fl^ media control condition. MOG_35-55_-reactivated cells were analyzed by ELISA for (**G, M**) IFNγ and (**H, N**) IL-17 secretion, and flow cytometry for (**I, O**) Tbet^+^ IFNγ^+^ Th1, (**J, P**) RORγt^+^IL-17^+^ Th17 and (**K, Q**) Tbet^+^IL-17^+^ cell populations. Data are pooled from four independent experiments, n = 6-10 mice. Mann-Whitney was performed for EAE score analysis; and One-way ANOVA, followed by Dunnett’s multiple comparison test. *p<0.05, **p<0.01, ***p<0.001, ****p<0.0001.

Since Th17 differentiation is abrogated in the absence of PRMT5, we expected this defect to be evident in peripheral T cell responses in the spleen. MOG-specific T cell proliferation (**Fig. 6L**), Th1 (**Fig. 6M, O**) and Th17 (**Fig. 6N, P**) responses, including the highly pathogenic Tbet^+^IL-17^+^ Th17 cells (**Fig. 6Q**), were robustly reduced among splenocytes. Similar abrogation of EAE disease and loss of Th1 and Th17 responses was observed in T-PRMT5^Δ/Δ^ mouse model where all T cells lack PRMT5 (**Supplemental Fig. 6**). Overall, these data reveal that Th-specific PRMT5 deficiency leads to powerful suppression of MOG-specific pathogenic Th17 and Th1 cell responses, demonstrating that PRMT5 expression in CD4^+^ T cells is necessary to drive pathogenic Th cell-mediated EAE autoimmunity.

## Discussion

Here we report that expression of the arginine methyltransferase PRMT5 sets the stage for Th17 differentiation, by promoting expression of enzymes in the cholesterol biosynthesis pathway that produce ROR-γt agonists. Consequently, PRMT5 in the Th cell compartment was absolutely required for the development of EAE. PRMT5 also modulates the thymic development and peripheral homeostasis stages of CD4 Th cells. We also observe robust T cell-intrinsic defects in iNK T and CD8 T cell development or maintenance, respectively, in the absence of PRMT5. Overall, the PRMT5^Δ/Δ^ mouse models reveal crucial roles of PRMT5 during thymic T cell development, peripheral homeostasis, Th cell polarization and T cell-mediated autoimmune disease.

The post-translational modifier, PRMT5, is essential for life (23) and has been implicated in stem cell function and development (3, 24, 25). PRMT5 plays crucial roles in carcinogenesis (26, 27) and autoimmune pathogenesis (6). Due to the well-established links between PRMT5 and proliferation, it was conceivable that PRMT5 deletion during thymocyte development would significantly impair thymocyte development. Indeed, loss of DP, CD4 SP and Treg cells is observed in the T-PRMT5^Δ/Δ^ model, consistent with CD4-driven cre mediated deletion at the DP stage. Thymic defects were not observed in another report of a constitutive CD4-cre driven PRMT5 KO mouse (8). This difference may be due to the deletion of all protein-coding isoforms in our KO model, compared to a KO of only the long isoform in the Inoue model (8). Alternatively, it may be mediated by the ubiquitous *Prmt5* heterozygous deficient background used in that model (8). The second possibility is highly likely, as we observed thymic defects in both PRMT5 homozygous and heterozygote mice in comparison to both PRMT5^fl/fl^ and CD4-cre controls (**Fig. 2**). While DP and CD4 SP numbers were decreased, their relative distribution remained stable, suggesting a defect mediated by impaired expansion. In contrast, CD8 SP numbers were normal, consistent with the higher proliferation that DP and CD4 SP cells vs. CD8 SP undergo during thymic development (28). Finally, as in the previous report (8), a striking defect was observed in iNK T cells, again consistent with the reported high proliferative needs of iNK T cells during thymic development (29).

Beyond thymic-level defects, we also observed reduced numbers of CD4, CD8 and NK T cells in the periphery, consistent with the observations in the Inoue pan-T cell PRMT5 KO model (8). The peripheral CD4 T cell loss likely derived from reduced entry into the peripheral compartment from the thymus and inability to homeostatically proliferate in response to IL-7 signaling, as previously proposed (8). The striking CD8 peripheral loss instead appears to exclusively stem from inability to survive and expand in the peripheral compartment, as thymic CD8 T cell numbers are normal. Finally, T cell proliferation in response to antigenic stimuli was abrogated in the absence of PRMT5. These data demonstrate that robust inhibition of T cell proliferation with PRMT5 inhibitors (6) is not due to off-target effects but indeed due to PRMT5 loss-of-function.

While multiple mechanisms tightly control PRMT5 expression in non-transformed T cells (5), uncontrolled PRMT5 expression and proliferation is a hallmark of hematologic and solid cancers (30-34). The driver role of PRMT5 in cancer has led to efforts to develop PRMT5 selective inhibitors as a therapy for solid and hematologic cancers, currently being tested for safety and efficacy (Clinicaltrials.gov, registration ID: NCT03573310, NCT03854227, NCT02783300, NCT03614728). Homozygous, but not heterozygote, PRMT5 deficiency early in bone marrow hematopoiesis results in severe bone marrow aplasia and eventual death (13). This phenotype has informed safety evaluation studies during clinical trials. We now show that complete PRMT5 deletion plays crucial roles in T cell development and biology, with robust suppression of Th17 and Th1 responses generally involved in pathogen protective immunity. It is important to note that more subtle or absent immune effects were observed in heterozygote Th PRMT5 KO mice, suggesting that careful PRMT5 inhibitor dosing could be the key to avoiding unintended effects. Such considerations will also be important when considering PRMT5 inhibitors for T cell-mediated diseases, albeit many of the T cell effects we here describe may instead serve a therapeutic role. Overall, it will be important to carefully monitor potential immune effects in PRMT5 inhibitor-based therapeutic approaches.

Our data show that PRMT5 promotes a cholesterol pathway metabolic switch, leading to expression of multiple biosynthetic genes in the cholesterol biosynthesis pathway. PRMT5 can promote gene expression in cancer cells by enhancing constitutive and alternative splicing (35, 36). A similar mechanism may promote cholesterol biosynthesis gene expression here. Several cholesterol pathway intermediates, including desmosterol, serve as agonists of RORγt and are required for Th17 differentiation (20-22). Supplementation with the cholesterol intermediate desmosterol restored Th17 differentiation in iCD4-PRMT5^Δ/Δ^ T cells, demonstrating that PRMT5 drives Th17 cell differentiation by promoting cholesterol biosynthesis. It is important to note that PRMT5 promotes cholesterol biosynthesis enzyme expression even in the absence of Th17-inducing cytokines (**Fig. 5A-C**). This is consistent with the idea that PRMT5 expression at the Th0 stage pushes Th17 differentiation via production of ROR-γt agonistic cholesterol pathway intermediates. These findings fit with observations in statin-treated Multiple Sclerosis patients. Statin drugs inhibit the HMG-CoA reductase rate-limiting step in the cholesterol biosynthesis pathway (37) and substantially reduce disability scores and atrophy in MS patients (38, 39). It has been unclear what is the mechanism behind these effects, but causal modeling analyses suggest an effect independent of peripheral cholesterol levels (40). Instead, these analyses implicate upstream intermediate metabolites of the cholesterol biosynthesis pathway (41) such as those promoted by PRMT5. Overall, the links between PRMT5, cholesterol intermediates pathway and pathogenic Th17 responses have therapeutic and biomarker implications in Multiple Sclerosis.

We found that PRMT5 expression in peripheral CD4^+^ T cells is an essential driver for the development of EAE disease. While pan-T cell T-PRMT5^Δ/Δ^ mice were completely resistant to EAE, this effect could be due to these mice lacking a full CD4, CD8 and iNK T cell compartments. Using the inducible peripheral CD4 Th-cell specific PRMT5 KO model (iCD4-PRMT5^Δ/Δ^), we confirmed that PRMT5 in CD4^+^ Th cells is required for EAE development. We observed strongly reduced proliferative, Th17 and Th1 recall responses in these mice, as well as severely reduced numbers of memory T cell infiltration in the CNS. The loss of Th17 responses was consistent with reduced Th17 differentiation. However, the Th1 cell response was also lost during EAE, in apparent contradiction to increased Th1 polarization observed (**Fig. 4B-E, K-N**). This result indicates that the EAE suppression in PRMT5 KO mice may be as a result of combined effects on polarization and expansion. Th1 polarization suppression may be compensated *in vivo* by enhanced expansion of Th1 cells during differentiation or reactivation stages. Indeed, PRMT5 inhibitors strongly suppress expansion of non-transformed Th1 memory T cell lines (6). Therefore, the overall impact of *in vivo* Th cell PRMT5 deficiency is therefore loss of both pathogenic Th1 and Th17 responses.

In summary, this study identifies a novel T-cell intrinsic role for PRMT5 as a regulator of T cell development, maintenance and Th17 cell differentiation and defines the metabolic switch mechanism by which PRMT5 controls Th17 differentiation. This metabolic switch consists of promotion of cholesterol biosynthesis after CD4^+^ Th cell activation towards ROR-γ agonist generation and activation of the Th17 program. The role of PRMT5 in mediating the pathogenic Th1 and Th17 cell responses identifies PRMT5 as a promising therapeutic target in MS and other Th17-mediated inflammatory diseases. Nonetheless, the crucial role of PRMT5 in T cell biology advises caution when using PRMT5 inhibitors as immunotherapies for autoimmune inflammatory disease or cancer.

## Materials and Methods

### Mice

All mice were bred and maintained under protocol #2013A00000151-R1. Sperm carrying the Prmt5^tm2c(EUCOMM)wtsi^ mutation was acquired from the Wellcome Trust Sanger Institute (Cambridgeshire, UK). A Prmt5^tm2c(EUCOMM)wtsi^ founder was obtained via *in vitro* fertilization at the OSU Comprehensive Cancer Center Animal Modeling Core (supported in part by grant P30 CA016058). Homozygous Prmt5^tm2c(EUCOMM)wtsi^ mice were bred with B6(129X1)-Tg(Cd4-Cre/ERT2)11Gnri/J or Tg(Cd4-cre)1Cwi CD4Cre mice from Jackson (Bar Harbor, Maine).

### In vivo tamoxifen treatment

Tamoxifen (T5648) was solubilized at 40mg/ml in corn oil by shaking at 37°C overnight. iCD4-PRMT5^fl/fl^ mice were given 300mg/kg tamoxifen (150ul per 20g mouse) daily for 3-5 days by oral gavage. Seven days after starting treatment, mice were euthanized or utilized for experiments. For extended tamoxifen treatment, mice were treated daily for 5 days every other week for a total of three sets of tamoxifen treatment. Mice were collected 3 days after the last day of tamoxifen treatment.

### T cell in vitro assays

Total CD4^+^ T cells were isolated using a CD4^+^ T cell isolation kit (Miltenyi Biotec 130-104-454, or Stem Cell Technologies #19852) and an autoMACS Pro (Miltenyi Biotec) or EasyEights magnet (Stem Cell). CD3^+^ (#19851), CD8^+^ (#19853), and/or naive CD4^+^ T cells (#19765) were isolated using Stem Cell EasyEights Magnet. For naive T cell differentiation experiments, naive T cells were differentiated in the presence of Th1 (IL-12, IL-2, anti-IL-4), Th2 (IL-4, IL-2, anti-IL-12, anti-IFNγ), Th17 (TGFβ, IL-6), or Treg (TGFβ, all-trans retinoic acid, IL-2) polarizing conditions. Isolated T cells were activated with 5µg/ml coated anti-CD3 and 2µg/ml soluble anti-CD28. T cells from iCD4-PRMT5^fl/fl^ mice were supplemented with 2µg/ml 4-hydroxytamoxifen (4-OHT; H7904) during *in vitro* cell culture.

### Flow cytometry

Thymocytes, splenocytes, lymph node cells, and bone marrow were taken directly *ex vivo* for flow cytometric analyses. MOG_35-55_-restimulated splenocytes and lymph node cells were treated with PMA/ionomycin and GolgiStop (BD) for 4 hours prior to collection for staining. Cells were treated with Fc region blockade and then incubated with surface stain markers, including B220 (BD 553087, clone RA3-6B2), CD3 (Biolegend 100334, clone 145-2C11), CD4 (Biolegend 100531 or eBioscience 12-0042-85, clone RM4-5), CD25 (Invitrogen RM6017 or eBioscience 45-0251-82, clone PC61 5.3), CD44 (eBioscience 48-0441-82 or 25-0441-82, clone IM7), CD62L (Biolegend 104426 or 104411, clone MEL-14), and CD8 (BD 560182, clone 53-6.7) for 15 minutes at 4 degrees. Cells were then fixed with eBioscience Fixation/Permeabilization buffer (eBioscience 00-5523-00), or BD Biosciences Fixation buffer (BD 554715). Intracellular staining with FoxP3 (eBioscience 17-5773-82, clone FJK-16s), T-bet (Biolegend 644810 or 644808, clone 4B10), IL-17 (Biolegend 506916, clone TC11-1810.1), IFN-γ (Biolegend 505830, clone XMG1.2), or RORγt (eBioscience 12-698880, clone AFKJS-9) was performed for 30-45 minutes at 4°C. Flow cytometry was run on a FACSCalibur with DxP multicolor upgrades (Cytek) and analysis was performed using FlowJo.

### ^3^H-Thymidine Proliferation assay

Isolated CD4^+^ T cells were plated at a density of 100,000-125,000 cells per well in a 96-well plate. After 48 hours of culture, cells were given 1µCi of tritiated (^3^H)-thymidine. 16 hours later, cells were harvested onto a Unifilter-96 plate (PerkinElmer 6005174). Scintillation fluid was added and counts per minute (CPM) were measured by a scintillation counter (Beckman Coulter).

### Western blot

Activated primary CD4^+^ T cells were collected directly *ex vivo* or activated and collected at indicated time points and cell pellets were frozen at −80°C. Cell pellets were lysed in RIPA buffer (10 mM Tris pH 7.4, 150 mM NaCl, 1% Triton X-100, 0.1% SDS, 1% deoxycholate) containing protease and phosphatase inhibitors (ThermoFisher). 5-10µg of protein were run on a 14% SDS-PAGE gel and transferred onto PVDF membrane. Blots were blocked with Odyssey Blocking Buffer (LICOR) and primary antibodies were probed for 3 hours at room temperature. Primary antibodies used include PRMT5 (ab31751, 1:1000), PRMT1 (CST #2449, 1:500), MYC (CST #9402, 1:500), Cyclin D1 (ab134175, 1:250), SYM10 (Millipore 07-412, 1:300), H4R3me2s (ab5823, 1:250), and B-actin (Sigma A1978, 1:20,000). Secondary antibodies anti-mouse 680RD and/or anti-rabbit 800CW (LICOR) were then used at 1:20,000. Blots were imaged on an Odyssey CLx (LICOR) and quantification of protein was performed using ImageStudio software.

### RNA sequencing

Isolated CD4^+^ T cells from PRMT5^fl/fl^ and iCD4-PRMT5^Δ/Δ^ mice (n=3 pooled mice/sample and n=3 samples per group) were stabilized in RNAlater (ThermoFisher AM7020) until RNA isolation. When all samples were ready for RNA isolation, the cell suspension was diluted in a 1:1 volume with 1xPBS prior to lysis with TRIzol (ThermoFisher 15596018). RNA isolation was performed with the Direct-zol RNA Miniprep (Zymo Research R2052) according to the manufacturer’s instructions. 1 ng of total RNA was used for quality control (QC), library preparation and RNA sequencing. The quality of total RNA was evaluated using Agilent 2100 Bioanalyzer and RNA Nano chip (Agilent Technologies, CA) and only samples with an RNA Integrity Number of ≥ 7.7 were sequenced.

RNA sequencing was performed by the Genomic Services Laboratory of the Abigail Wexner Research Institute at Nationwide Children’s Hospital, Columbus, Ohio. RNA-seq libraries were prepared using Illumina’s TruSeq Stranded protocol. In summary, ribosomal RNA (rRNA) was removed from 250 ng of total RNA with Ribo-zero Human/Mouse/Rat Gold kit. The kit depletes samples of both cytoplasmic and mitochondrial rRNA using biotinylated, target-specific oligos combined with rRNA removal beads. Following rRNA removal, mRNA is fragmented using divalent cations under elevated temperature and converted into ds cDNA. Incorporation of dUTP in place of dTTP during second strand synthesis inhibits the amplification of the second strand. The subsequent addition of a single ‘A’ base allows for ligation of dual unique tagging sequences. Adaptor-ligated cDNA was amplified by limit-cycle PCR. Quality of libraries were determined via Agilent 4200 Tapestation using a High Sensitivity D1000 ScreenTape Assay kit, and quantified by KAPA qPCR (KAPA BioSystems). Approximately 60-80 million paired-end 150 bp sequence reads were generated for each library on the Illumina HiSeq 4000 platform.

For analysis, low-quality reads (q<10), and adaptor sequences were eliminated from raw reads using BBDuk version 37.64 (DOE Joint Genome Institute). Each sample was aligned to the GRCm38.p3 assembly of the *Mus musculus* reference from NCBI using version 2.6.0c of the RNA-Seq aligner STAR (42). Features were identified from the GFF file that came with the assembly from Gencode (Release M19) and feature coverage counts were calculated using feature Counts (43). Differentially expressed features were calculated using DESeq2 (Bioconductor Release 3.9).

### Real time PCR

To evaluate mRNA expression, 300 ng-500 ng of RNA were reverse transcribed using oligo d(T) or random primers and Superscript III (ThermoFisher18080051) according to manufacturer’s instructions; TaqMan quantitative real-time PCR was performed using mouse *Prmt5* (Mm00550472_m1), mouse *Hprt* (Mm0044968_m1), mouse *Cyp51a* (Mm00490968_m1), mouse *Tm7sf2* (Mn0123354_g1), mouse *Lss* (Mm00461312_m1), mouse *Nsdhl* (Mm00477897_m1), mouse *Acat2* (Mm00782408_s1) and mouse *Hmgcs1* (Mm01304569_m1) as previously described (ThermoFisher). Samples were run on Quant Studio 5 (ThermoFisher). An initial denaturation step at 95°C for 10 min was followed by 40 cycles of denaturation at 95°C for 15 s and primer annealing/extension at 60°C for 60 s. Results were analysed using the comparative Ct method after ensuring comparable amplification efficiencies for test and housekeeping transcripts.

### Experimental autoimmune encephalomyelitis

To induce EAE, mice were immunized with myelin oligodendrocyte glycoprotein peptide (MOG_35-55_; CS Bio) and CFA (Difco) emulsion, as previously described (6). Mice were monitored for disease every day and scored blinded. At the indicated time points, mice were euthanized by injection with 20 mg/ml ketamine and 4 mg/ml xylazine (150 µl/20 g mouse) and perfused with PBS. Spleens, brains, and spinal cords were collected from representative mice and processed for *in vitro* studies. To isolate brain and spinal cord mononuclear cells, brains and spinal cords were processed through a 70-μm strainer and separated by a 70–30% isotonic Percoll gradient. Spleens were processed through a 70-μm strainer and red blood cells were lysed by incubating for 1 minute in hypotonic solution. Splenocytes and infiltrating CNS cells were used for *ex vivo* flow cytometry or reactivated with MOG antigen as indicated.

### Statistical analysis

Statistical analyses were performed using GraphPad Prism unless otherwise stated. One-way ANOVA and Student’s *t* tests were used as appropriate, and as indicated. Mann-Whitney test was performed for EAE score data analysis.

### Study approval

All animal studies were performed after appropriate institutional eIACUC approval, under protocol #2013A00000151-R1.

## Acknowledgements

We would like to thank the Genomic Services Laboratory of the Abigail Wexner Research Institute at Nationwide Children’s Hospital for their help with RNAseq. We thank Amy Wetzel, Shireen Woodiga, Anthony Miller and Saranga Wijeratne of the Genomic Services Laboratory at the Abigail Wexner Research Institute at Nationwide Children’s Hospital, Columbus, Ohio for their help with sample QC, library preparation, RNA sequencing and analysis of data. We also thank Dr. Ning Quan, Dr. Gene Oltz and Dr. Philip Tsichlis for their insightful discussions and comments.

## Author contributions

MGdA, LW and SS contributed to conceptual experimental design. LW, SS, CE, SAA, AK and JNM performed experiments and analyzed data. MGdA, LW and SS interpreted data and wrote the manuscript. All authors thoroughly reviewed the manuscript.

## Materials and correspondence

Correspondence and material requests should be addressed to Dr. Mireia Guerau-de-Arellano. School of Health and Rehabilitation Sciences. Division of Medical Laboratory Science. 453 W 10^th^ Ave. Columbus Oh 43210. Atwell Hall 235A.

