## Supplemental Figures 1-6 for "PRMT5 in T helper lymphocytes is essential for cholesterol biosynthesis-mediated Th17 responses and autoimmunity"

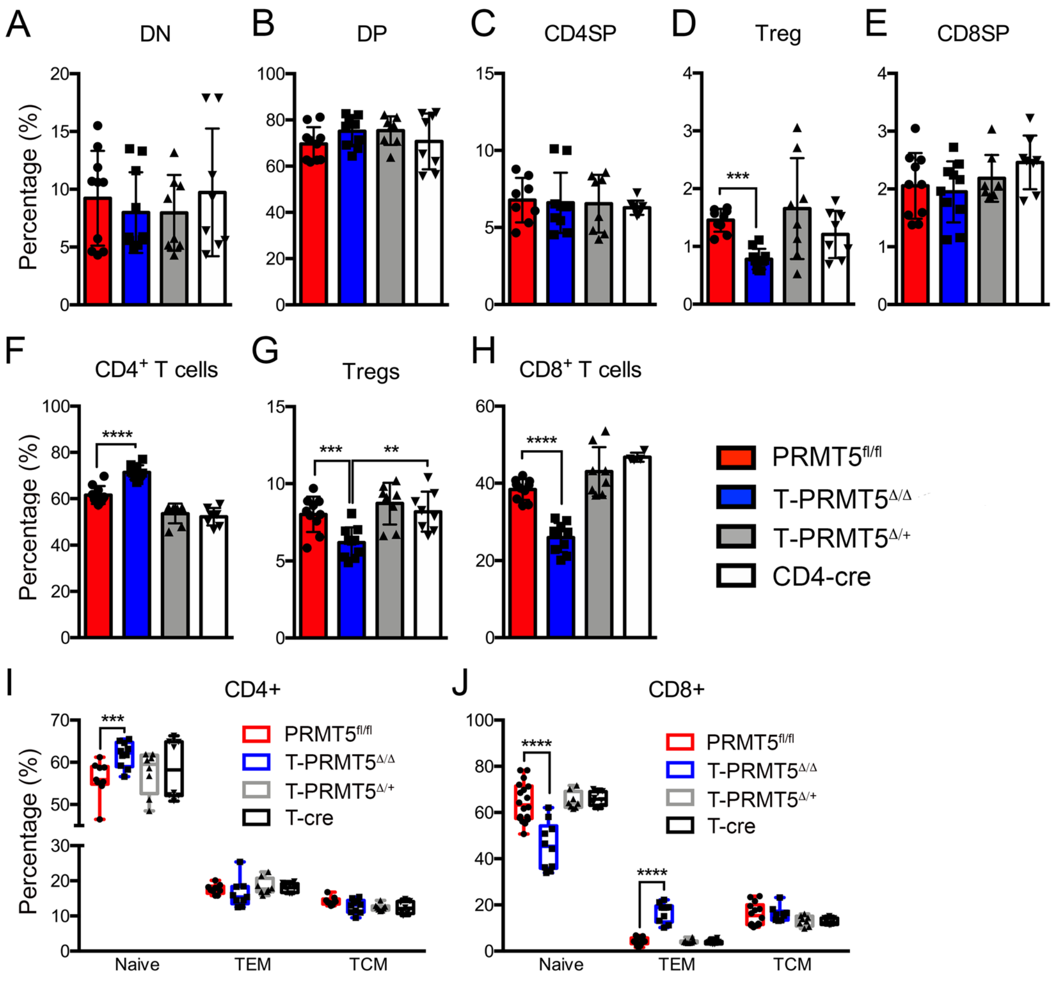


**Supplemental Figure 1. Impact of T-PRMT5$^{\Delta/\Delta}$ on thymic and peripheral immune cell frequencies.**

(**A-E**) Thymocytes from T-PRMT5$^{\Delta/\Delta}$ and appropriate control mice were analyzed by flow cytometry for frequencies of (**A**) DN, (**B**) DP, (**C**) CD4SP, (**D**) Treg, and (**E**) CD8SP cell populations. (**F-H**) Splenocytes were analyzed by flow cytometry for percentages of (**F**) CD4$^{+}$, (**G**) Tregs, (**H**) CD8$^{+}$, (**I**) $T_{EM}$, $T_{CM}$, and naive CD4$^{+}$ T cell subsets and (**J**) $T_{EM}$, $T_{CM}$, and naive CD8$^{+}$ T cell subsets. Data are pooled from 2 independent experiments (shown n=4-5). One-way ANOVA, followed by Dunnett's multiple comparison test. *p<0.05, **p<0.01, ***p<0.001, ****p<0.0001.

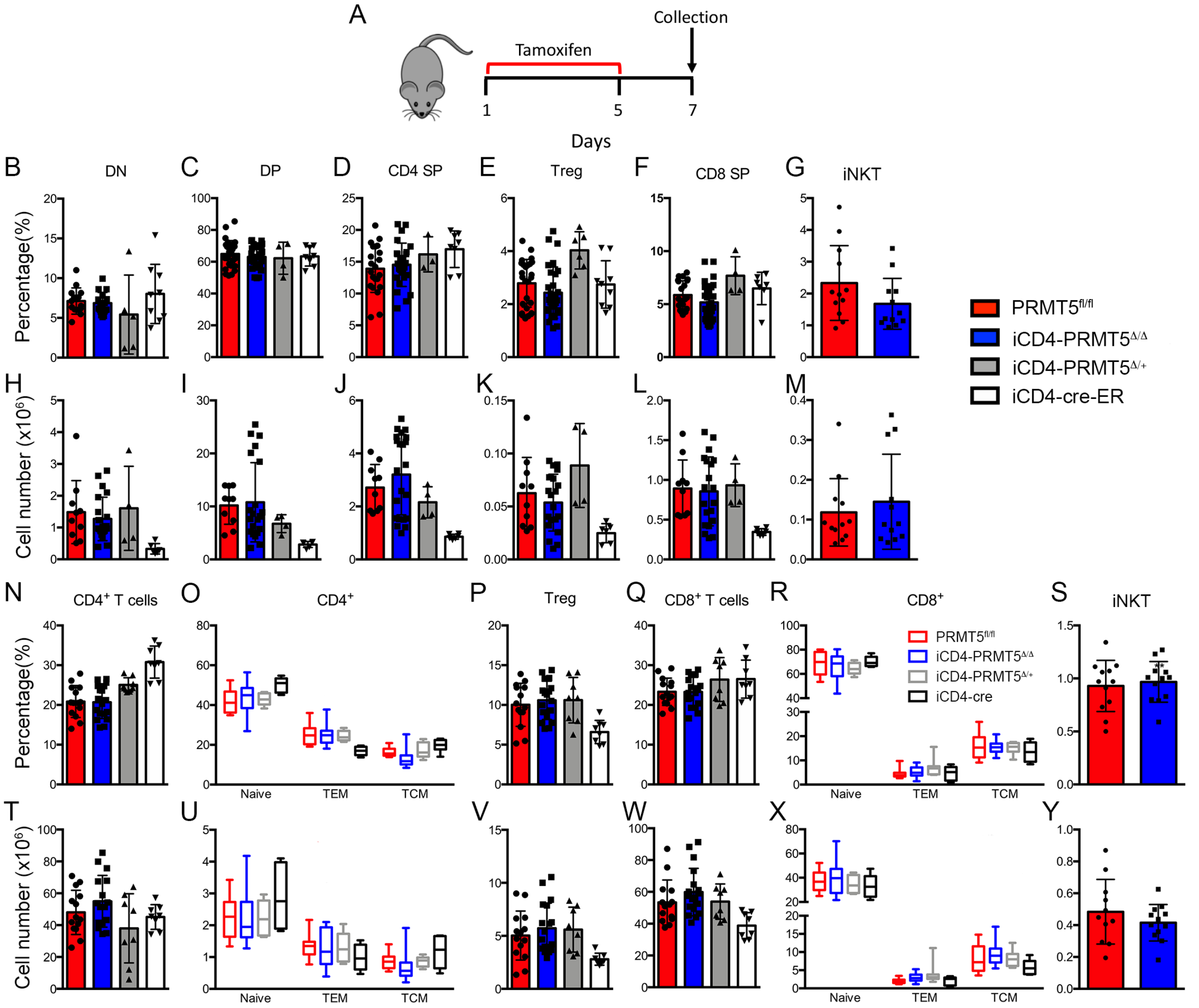


**Supplemental Figure 2. Acute PRMT5 knockout in CD4$^{+}$ T cells does not affect thymic development or peripheral immune cell compartments.**

(**A**) Schematic of tamoxifen treatment experimental design and collection for direct ex-vivo flow cytometry analyses. (**B-M**) Thymocytes from iCD4-PRMT5$^{\Delta/\Delta}$ and appropriate control mice treated with tamoxifen for one week to induce acute peripheral CD4 T cell PRMT5 deletion were analyzed by flow cytometry. (**B-G**) Frequency and (**H-M**) cell number of thymic (**B, H**) CD4$^{-}$ CD8$^{-}$ DN, (**C, I**) CD4$^{+}$CD8$^{+}$ DP, (**D, J**) CD4SP, (**E, K**) Treg, (**F, L**) CD8SP and (**G, M**) iNK T cell populations. Splenocytes were analyzed by flow cytometry for (**N-S**) percentages and (**T-Y**) cell numbers of (**N, T**) CD4$^{+}$, (**O, U**) CD4$^{+}$ $T_{EM}$, $T_{CM}$, and naive, (**P, V**) Tregs, (**Q, W**) CD8$^{+}$, (**R, X**) CD8$^{+}$ $T_{EM}$, $T_{CM}$, and naive and (**S, Y**) iNK T cell populations. Data are pooled from at least 4 independent experiments (shown n=6-8). One-way ANOVA, followed by Dunnett's multiple comparison test. *p<0.05, **p<0.01, ***p<0.001, ****p<0.0001.

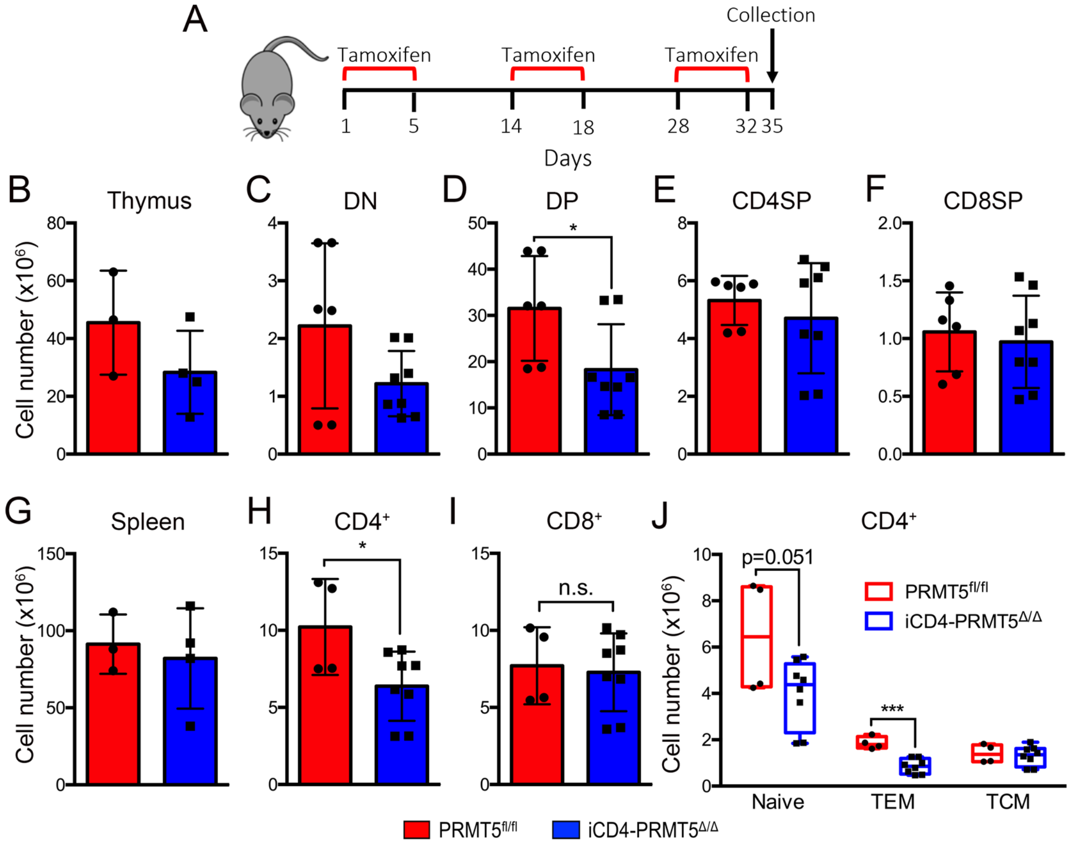


**Supplemental Figure 3. Impact of extended *Prmt5* deficiency in iCD4-PRMT5$^{\Delta/\Delta}$ mice.**
(**A**) Schematic of experimental design for long-term tamoxifen treatment of iCD4-PRMT5$^{\Delta/\Delta}$ and PRMT5$^{fl/fl}$ mice and collection for direct ex-vivo flow cytometry analyses. (**B-F**) Thymi were processed and (**B**) total cell numbers were counted. Flow cytometric analysis was performed and cell numbers were calculated for (**C**) DN, (**D**) DP, (**E**) CD4SP and (**F**) CD8SP compartments. (**G-J**) Splenocytes were isolated and (**G**) total cell numbers were counted. Flow cytometric analysis was performed and cell numbers were calculated for (**H**) CD4$^+$, (**I**) CD8$^+$ and (**J**) CD4$^+$ $T_{EM}$, $T_{CM}$, and naive, cell compartments. Data are pooled from 3-4 independent mice. Student's *t* test. *p<0.05, **p<0.01, ***p<0.001, ****p<0.0001.

A

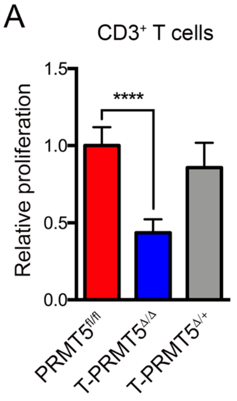


**Supplemental Figure 4. Impact of *Prmt5* deficiency on $CD3^+$ T cell proliferation.**
(**A**) $CD3^+$ T cells were isolated from T-PRMT5$^{\Delta/\Delta}$ and control mice and activated on anti-CD3/CD28 for 48 hours. Proliferation was monitored by $^{3}$H-thymidine incorporation. Data are pooled from 2 independent experiments (shown n = 3-4). One-way ANOVA, followed by Dunnett's multiple comparison test. *p<0.05, **p<0.01, ***p<0.001, ****p<0.0001.

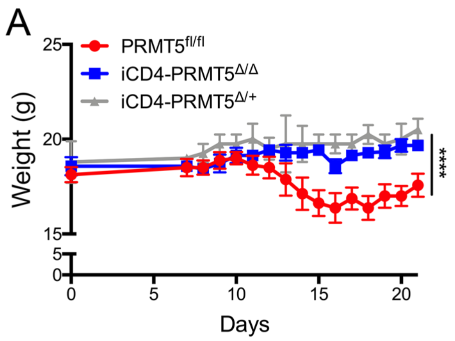


**Supplemental Figure 5. T cell specific *Prmt5* deficiency prevents induction of EAE autoimmunity** (**A**) iCD4-PRMT5$^{\Delta/\Delta}$ and appropriate control mice were immunized with CFA/MOG and weights of EAE mice were monitored daily. Data are pooled from four independent experiments (shown n = 6-10). Mann-Whitney was performed for EAE score analysis; *p<0.05, **p<0.01, ***p<0.001, ****p<0.0001.

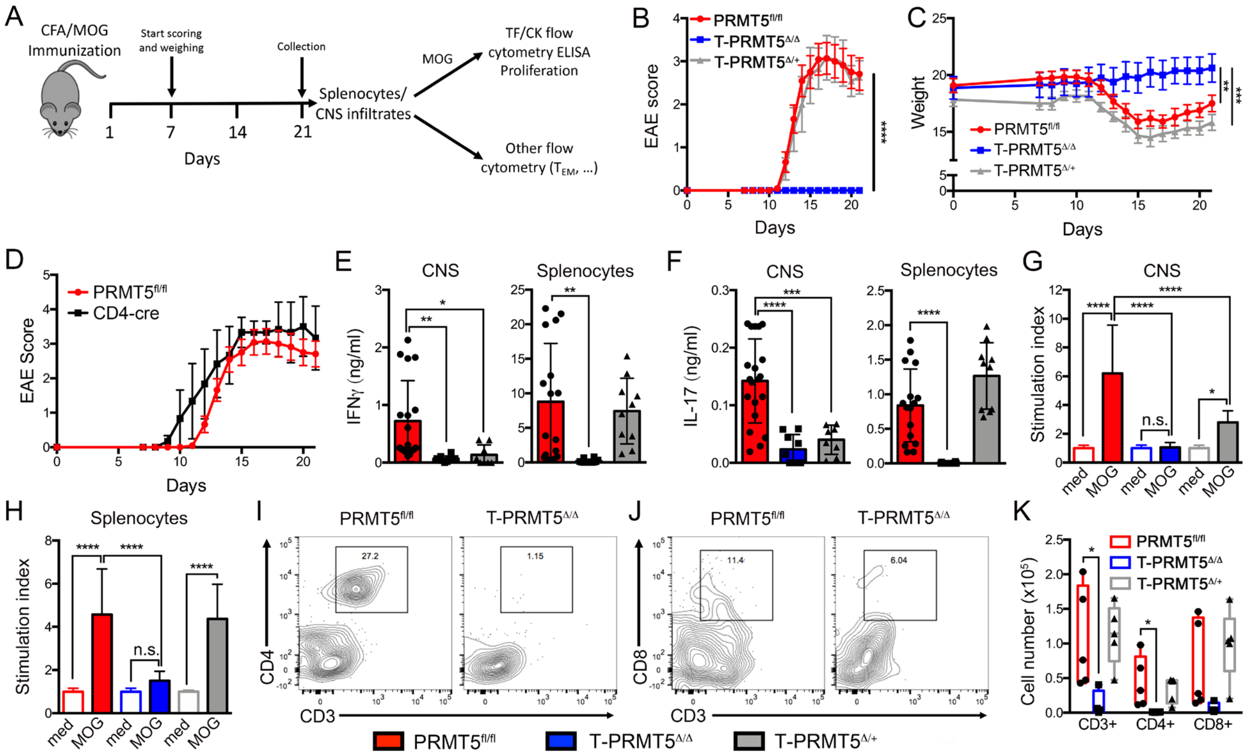


**Supplemental Figure 6. T cell specific *Prmt5* deficiency prevents induction of EAE autoimmunity.**

(**A**) Schematic of EAE experimental design and downstream analyses. (**B**) EAE score in T-PRMT5$^{\Delta/\Delta}$ and indicated controls after MOG$_{35\text{-}55}$/CFA immunization. (**C**) Weights of EAE mice were monitored daily. (**D**) Scores of PRMT5$^{fl/fl}$ (n=10) and additional CD4-cre control (n=3) mice were monitored daily. (**E-H**) Splenocytes and infiltrating CNS cells were isolated at day 21 after MOG$_{35\text{-}55}$/CFA immunization, and reactivated with MOG to measure (**E**) IFNγ and (**F**) IL-17 production by ELISA and (**G-H**) proliferation by $^{3}$H-thymidine incorporation. (**I-K**) Flow cytometric analysis of *ex vivo* infiltrating CNS cells quantifying (**K**) CD3$^{+}$, (**I, K**) CD3$^{+}$CD4$^{+}$, and (**J, K**) CD3$^{+}$CD8$^{+}$ populations at day 21. Data are pooled from four independent experiments, n=6-10 mice. Mann-Whitney was performed for EAE score analysis; For other analyses, one-way ANOVA, followed by Dunnett's multiple comparison test was used. *p<0.05, **p<0.01, ***p<0.001, ****p<0.0001.
